# Cell size regulation in bacteria

**DOI:** 10.1101/004457

**Authors:** Ariel Amir

## Abstract

Various bacteria such as the canonical gram negative *Escherichia coli* or the well-studied gram positive *Bacillus subtilis* divide symmetrically after they approximately double their volume. Their size at division is not constant, but is typically distributed over a narrow range. Here, we propose an analytically tractable model for cell size control, and calculate the cell size and inter-division time distributions, and the correlations between these variables. We suggest ways of extracting the model parameters from experimental data, and show that existing data for *E. coli* supports *partial* size control, and a particular explanation: a cell attempts to add a *constant volume* from the time of initiation of DNA replication to the next initiation event. This hypothesis accounts for the experimentally observed correlations between mother and daughter cells as well as the exponential dependence of size on growth rate.

PACS numbers: 87.17.Ee, 87.17.Aa, 87.10.Mn, 87.81Tt

---

Microorganisms such as bacteria come in a diverse set of shapes and sizes. Nonetheless, individual strains have remarkably reproducible shapes, and a narrow distribution of sizes [1–4]. Many bacteria, such as *E. coli*, are rod-shaped, and during their exponential growth phase they elongate while maintaining a constant diameter. After approximately doubling their length (as well as mass and volume), and completing DNA replication for their offspring, they divide symmetrically into two approximately identical daughter cells. In spite of decades of research, we still do not have a good understanding of how cells regulate their shape, both mechanically (i.e., what is the biophysical feedback necessary to achieve a rod-shape cell? [5]) and dimensionally: the coefficient of variation (standard deviation:mean, CV) can be as low as 0.1 for bacteria [2]. Bacteria are also remarkable in their ability to have a generation time that is shorter than the time it takes them to replicate DNA: doubling time τ*_d_* for *E. coli* in rich media at 37°C is about 20 mins, while T*_r_*≈ 60 mins are needed from initiation of DNA replication to cell division. This apparent paradox is explained by the existence of *multiple replication forks* : in these situations, a cell will already start replicating DNA for its 4 granddaughters (or 8 great-granddaughters), in order for the replication to complete in time.

Many models for cell size regulation exist in the literature [1, 2, 6–14]. Different strategies will yield particular cell size and inter-division time distributions, as well as distinct correlations. Hence, it is important to understand the connection between different regulation models and the resulting distributions and correlations. Moreover, there are two seemingly contradictory results in the literature: the first is the model by Donachie [15], which shows that the measured exponential dependence of bacterial size on growth rate [16] is consistent with initiation of DNA replication at a constant, growth-rate-independent volume per replication fork - suggesting a mechanistic picture in which a cell “knows” of its size and initiates replication when reaching a critical one. This model would imply that size at birth and division would not be correlated: since the time from initiation to division is constant [17], the size at division will be independent of the size at birth. However, experiments show that there are strong correlations between the two [18].

We will show here how these two results can be elegantly reconciled within a minimal model, which will be analytically tractable. We will suggest a mathematical framework which is able to capture and extend several existing models, and will use it to analyze the correlations and cell size distributions. We shall show that the aforementioned experimental data for E. coli supports a mechanism of cell size regulation in which the cell attempts to *add a constant volume from the event of initiation of DNA replication to the next initiation event* [19]. This model will be consistent with the results discussed in Ref. [15], predicting an exponential dependence of cell size on growth rate, but will also quantitatively account for the positive correlations between size at birth and division [18] and negative correlations between size at birth and inter-division time [14]. We will show that for size-additive noise the size distribution is Gaussian, while for time-additive (i.e., size-multiplicative) noise the resulting size distribution is log-normal - and hence right-skewed.

As shown in the Supplementary Information (SI), experimentally measured distributions are indeed skewed, and for this reason we focus on the analysis of time-additive noise in the main text and defer the size-additive case to the SI. The standard deviations of both size and interdivision time distributions are controlled by a *single* parameter.

The tools which we shall use will parallel those used when solving problems in statistical mechanics, in particular those involving Langevin equations [20]. Multiplicative noise and the log-normal distributions which emerge from our model also occur in other problems in physics, such as relaxations in glasses [21] and the modelling of financial markets [22, 23]. However, in contrast to most physical systems, negative feedback (i.e., control) is a necessary feature of biological systems, including the problem studied here.

## Exponential growth of a single cell and regulation models

The question of the mode of growth of a single bacterium has been a long standing problem, with linear and exponential growth the most common models considered [1, 2, 24]. Recent experiments show that individual cell volume grows exponentially, for various bacterial strains [3, 25–27]. In fact, if cells grow at a rate that is proportional to the amount of protein they contain [28, 29], as long as the protein concentration is constant, the cells will grow exponentially in mass and volume. We shall assume exponential growth of *volume* throughout this paper, *v*(*t*) ∝ *2^t/^*^τ*d*^, and neglect fluctuations in the growth rate. Furthermore, we will assume that cells divide precisely in half since experimental results [30] show that division occurs at the mid-cell to an excellent approximation.

Cells need a feedback mechanism that will control their size distribution. If cells grew for a constant time *t* = *τ_d_*, random fluctuations in the timing would make the size of the cells at division, *v_d_*, perform a random walk on the volume axis, and thus this mechanism does not control size. Another regulatory strategy is that of division at a critical mass, or of initiation of DNA replication at a critical size. These ideas are prevalent in the literature [1, 2], but we will show that existing experimental data for *E. coli* argues against them. We shall consider the following class of models: upon being born at a size *v_b_*, the cell would attempt to grow for a time *τ* (*v_b_*) such that its final volume at division is *v_d_* = *f*(*v_b_*). If the function *f*(*v_b_*) = *const*, we are back to the critical mass model. The constant time model can also be cast in this language: since the growth is exponential, attempting to grow for a prescribed, constant time *τ_d_* is the same as having *f*(*v_b_*) = 2*v_b_*. Another important model that has been suggested is the so-called “incremental model”, in which the cell attempts to add a constant volume *v*_0_ to its newborn size [31]. In this case:

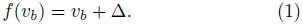

In the following, we suggest a method through which an arbitrary regulatory model described by a function *f*(*v_b_*) can be approximately solved, i.e., we can find all the involved distributions and correlation coefficients analytically, finding excellent agreement with the numerically exact solutions. We also provide methods to extract the model parameters from experimental data.

## The model

We assume that the cell attempts to divide at a volume *v_d_* = *f*(*v_b_*), as previously explained, by attempting to grow for the appropriate amount of time *t_a_* which is a function of *v_d_*. We assume that to this time is added a random noise *t_n_*, which we assume to be Gaussian. The magnitude of this noise will dictate the width of the resulting size and inter-division time distributions. Thus we have:

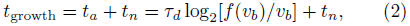

with *t_n_* assumed to be a random variable with: 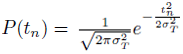. The model is similar to that discussed in Ref. [32], where the molecular mechanisms leading to the noise in budding yeast are studied.

We will calculate the inter-division time and volume distributions. The key insight is that for noise that is not too large (equivalent to size distributions which are not too broad, i.e., with a small CV), it is the behavior of *f* (*v_b_*) around the average newborn size 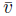 that is the most important. Therefore we can Taylor expand *f* (*v_b_*) around 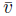:

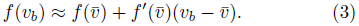

As an example, the incremental model has 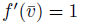 and 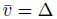, while the critical size model has 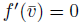.

Any two models that agree to lowest order, will result in similar distributions - provided the noise is not too large. We therefore choose to solve an equivalent model, that will be amenable to analytic treatment, and that can be viewed as an interpolation between the critical size model and the constant doubling time model. We choose:

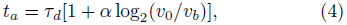

which corresponds to the regulatory function: 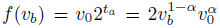. The case α = 0 corresponds to constant doubling time model (*f′*(*v*_0_) = 2), while *α* = 1 corresponds to the critical size model (*f′*(*v*_0_) = 0). Importantly, for *α* = 1/2 we have *f′*(*v*_0_) = 1, as does the incremental model: hence, using a target function like this gives results close to a perfect realization of the incremental mode. We shall show that the paramter *v*_0_ in Eq. (14) will be very close to the average newborn cell size 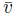.

## Solution of size and inter-division time distributions

We shall consider the case of *symmetric* division, relevant for many bacteria. For a newborn size *v*_*b*_, we have for the next newborn volume: 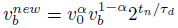.

Therefore:

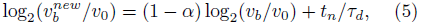

From stationarity of the stochastic process we know that 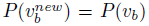. Since *t_n_* is a Gaussian variable, we find that log_2_(**v*_b_*) is also a Gaussian variable, and hence *P*(**v*_b_*) would be a *log-normal* distribution. If we denote the variance of log_2_(**v*_b_*/*v*_0_) by 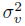, we have 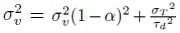, therefore the newborn size distribution is:

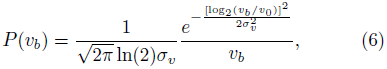

with

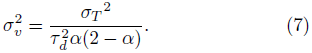

Note that the average cell size is 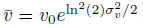; for realistic values of *σ_ν_* it will only be a few percent larger than *v*_0_. Similarly, the standard deviation of the size distribution will be approximately *σ_s_* ≈ ln(2)*σ_v_v*_0_, and the coefficient of variation is thus: *v*_*CV*_ ≈ ln(2)*σ_v_*. The skewness of the distribution is positive: γ_1_ ≈ 3ln(2)*σ_v_*, and provides a useful test of the assumption of a time-additive rather than size-additive noise, as we elaborate on in the SI.

We can now find the distribution of division times using: *t_d_* = *t_a_* + *t_n_*. Since **v*_b_* depends only on the noise of previous generations, *t_a_* is independent of *t_n_*, and since log_2_(**v*_b_*/*v*_0_) and *t*_*n*_ are Gaussian variables, the resulting inter-division time distribution is also Gaussian, and has a variance given by:

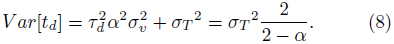

In the case a *α* → 0, we find that *σ_ν_* diverges (an extremely broad distribution of newborn sizes), but the inter-division time distribution is narrow: *Var*[*t_d_*] → *σ_T_*^2^, as should clearly be the case since there is no size feedback mechanism in this case. Note that for any positive value of α > 0 there will be a stationary size distribution, but for *α* = 0 there is no stationary distribution.

From Eq. (16) we find that the CV of the distribution of inter-division times is given by: 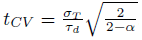. It is instructive to consider the dimensionless quantity:

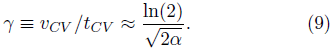

By constructing γ from the experimental distributions we can extract the value of α and find the form of size regulation utilized by the organism, if the division is symmetric. Later we shall show an additional, independent way of extracting *α*, which will be more robust against measurement noise since it will rely on correlations rather than the distribution widths.

Fig. 1 compares the numerically obtained size distribution for various values of α and the incremental model, with the result of Eq. (6), finding excellent agreement. In the SI we extend this comparison to various noise magnitudes. Our model captures the numerically exact solution very we11 and Eq. (14) provides a usefu1 too1 to capture a generic division strategy characterized by an arbitrary function *f*(**v*_b_*).

**FIG. 1.**
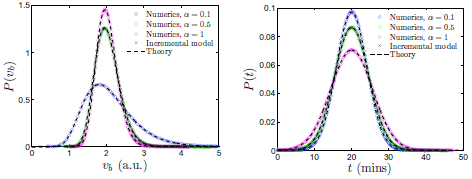
Comparison between the analytical results of the model for varying values of *α* (Eqs. (6–16)), and numerics. Choosing *α* = 1/2 provides an excellent approximation for the incremental model, as the effective size regulation of the two models agrees to lowest order. The parameters of the model are chosen according to their realistic values for *E. coli* growing at 37^o^: doubling time is *τ_d_* = 20 mins and *σ*_*T*_/τ*_d_* =0.2 [2]. For each case, the numerical distribution is extracted from a sequence of 10^7^ divisions.

## Extracting the parameters from experiments

Within the class of models proposed here, the value of *α* can be obtained by considering the correlations between size at birth and size at division. In the SI we show that the Pearson correlation coefficient between size at birth and division (equivalent to the mother:daughter correlation coefficient) equals:

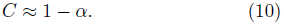

In particular, for the incrementa1 mode1 the correlation coefficient between mother and daughter cells should be 0.5.

Upon fixing the value of *α*, a *single* parameter, *σ_T_*, will determine the distributions of both size and division time, and the calculations performed here would allow one to scale both distributions using this single parameter. For the time-additive noise analyzed here, the model predicts an approximately log-normal newborn size distribution, given by Eq. (6), and a Gaussian inter-division time distribution, given by Eq. (16), whose standard deviation is larger than *σ_T_*. In the SI we show that for size-additive noise, one obtains a skewed time-distribution but a Gaussian size distribution - in contrast to what is observed experimentally [13]. Therefore, observing the distribution shape provides useful information regarding the source of the noise. Further experiments are needed to elucidate the molecular source of this multiplicative noise.

## Cell size control in E. coli

Experimentally, various correlation coefficients were measured for *E. coli* at slow growth conditions in Ref. [18], using the membrane elution technique. The correlation coefficient between new-born cell and daughter cell size was found to be *C* = 0.55, close to the theoretical 1/2 value expected for the incremental model. There was a strong correlation (0.8) between size at initiation of DNA replication and size at division, as we would expect from the assumption of exponential growth and that the time from initiation to division is constant [17]. Yet these observations appear to be in direct contradiction to the idea that initiation occurs at a critical size [15]. The key point is that Donachie’s analysis [15] shows that there is a critical size for initiation of DNA replication (independent of growth rate), *on average*. It is only from the fluctuations (i.e., correlations) that one can understand whether the underlying regulatory mechanism utilizes a critical size or integrates volume - as we shall propose is the case. Ref. [19] gives a simple biophysical implementation of the incremental model, which will reconcile these seemingly contradictory results and will realize a particular case of the class of model we proposed here: in this model, a protein *A* is forced to have a growth-rate-independent density throughout the cell using a negative feedback in its regulation, and a second protein *B* is produced whenever *A* is. In this way when cell volume grows (and only then), more *A* and *B* proteins are generated in an amount proportional to the change in *volume*. The hypothesis is that *B* proteins localize at their potential initiation site (which we will assume to be one of the replication origins), and only when their total *number* at each origin reaches a critical value does initiation of DNA replication occur, after which *B* is degraded. Note that two types of proteins are necessary, since in order to measure volume differences *A* must be spread throughout the cell, while *B* has to localize to measure an absolute number (rather than concentration). See Ref. [19] for further details.

In the SI we show that this model reduces to the incremental model, albeit with an effective Δ (see Eq. (1)) which strongly depends on the growth rate:

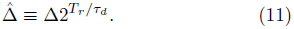

According to our results the average cell size will be Δ̂ - in agreement with the experimental results seeing precisely this exponential dependence of bacterial size on growth rate, with *T_r_* the exponent [16]. This model naturally accounts for the “quantization” of the cell critical mass at initiation at different growth rates [15], without necessitating the measurement of an absolute mass or volume. Moreover, it is plausible that the source of noise in adding the incremental volume will be due to “molecular noise” (number fluctuations of protein *B*), and would therefore be weakly dependent on growth rate. The same calculation which leads to Eq. (21) would suggest that *σ_T_* (the noise standard deviation) should depend on the growth rate in the same exponential way as Δ̂. This implies that the CV of size distributions should be weakly dependent on growth rate (see Eq. (7)), an expectation supported by Ref. [33].

Thus, we have shown that using our calculations and the interpretation in terms of the incremental model explains various experimental results. In fact, the model also makes precise predictions with regards to additional correlations: for example, it is possible to show that for the incremental model the size correlation coefficient between cells *N* generations apart is 2^−*N*^. Similarly, the model predicts a negative correlation of -1/2 between the size at birth and the inter-division time, see the SI for further details. This correlation coefficient was recently analyzed by Robert et al. [14], using data from two different experimental systems (with different growth rates), finding a correlation coefficient of -0.5 in both cases: exactly as predicted by our model. This gives a particularly simple and transparent interpretation to their analysis, and provide additional, strong support for the incremental model. Refs. [13 and 34] find similar negative correlations between newborn size and inter-division size, supporting our conclusion.

All of these provide additional support for the relevance of this model to cell size control in *E. coli*, and most likely to other organisms as well. It is possible, however, that alternative biophysical mechanisms may lead to the same correlations and size dependencies calculated here, and for this reason finding the underlying biological mechanism is important; In recent years, dnaA has been shown to have properties reminiscent of the biophysical model described here [9], where its active and inactive forms correspond to the roles of proteins *A* and *B* above - see the SI for further details.

## Discussion

In this work we suggested a phenomenological model which is able to describe partial size control within a broad class of control strategies, and interpolate between the case of constant time to division and division at a critical size, for both size-additive and time-additive noise. We are able to analytically calculate the size and inter-division time distributions for the case of symmetric division, relevant to various bacteria. For *E. coli*, we have shown that a simple biophysical model in which a constant volume is added from consequent events of initiation of DNA replication predicts: 1) Cell size depends exponentially on growth rate. 2) Cell size distributions are approximately log-normal. 3) The correlation coefficient between size at birth and division is approximately 1/2. 4) The correlation coefficient between size at birth and time to division is approximately -1/2. 5) The ratio of the CV of size and inter-division time distributions is approximately ln(2). The simplicity of a biophysical model which implements this idea [19] suggests that this may be a robust way of regulating cell size and coupling DNA replication and growth.

This interpretation in terms of the incremental model suggests an outstanding puzzle: can we underpin the precise molecular mechanism responsible for volume integration? Can the source of the noise in inter-division times be elucidated? Testing this model further in other microorganisms may yield important insights into cell size regulation, and in particular, it is intriguing to see if the same ideas are applicable to cell size control in higher organisms. Recently, size distributions in other microorganisms were shown to obey simple scaling laws [35], suggesting this to be a promising direction, and that the model discussed here may have a broader range of applicability.

## Acknowledgments

This research was supported by the Harvard Society of Fellows and the Milton Fund. We thank Sriram Chandrasekaran, Suckjoon Jun, Andrew W. Murray, Johan Paulsson, Ilya Soifer and Sattar Taheri-Araghi for useful discussions, and to Johan Paulsson for introducing the incremental model to us. We also thank the referees for useful suggestions.

## CELL SIZE REGULATION IN MICROORGANISMS - SUPPLEMENTARY INFORMATION

### Calculating the correlation coefficients

In the following we calculate the Pearson correlation coefficients between various variables of interest, for the model with an arbitrary value of *α* characterizing the size control strategy.

#### Mother size:daughter size correlation coefficient

For a narrow size distribution (such as the biologically relevant one), performing a linear regression analysis between the size at birth and the size at division would not be very different than doing the regression between *x* = log_2_(**v*_b_*/*v*_0_) and *y* = log_2_(**v*_d_*/*v*_0_) (for the data corresponding to Fig. 1, for example, the difference between the two methods is less than one percent). Within our model, the value of the dimensionless slope *β* can be readily found from Eq. (3) of the main text: since the noise is uncorrelated with the random variable *x*, we have:

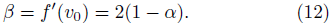

(i.e., it is 1 for the incremental model). For symmetric divisions, the slope of the linear regression between a newborn cell and the size of the daughter cell immediately after division would thus equal 1 − *α*. The coefficient of correlation between the newborn cell and the newborn daughter cell is:

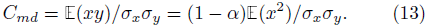

From stationarity, we know that the distribution of **v*_b_* and of 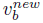 are identical, hence *σ_y_* = *σ_x_*, and: *C* = 1 − *α*. Therefore, for the incremental model the correlation coefficient between mother and daughter cells should be 0.5.

#### Newborn Size: Generation time correlation coefficient

From Eq.(4) of the main text:

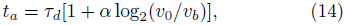

Hence, we expect a negative correlation between log(newborn size) and inter-division time, with slope −*α*. Such a negative correlation was recently observed [1]. Let us calculate the Pearson correlation coefficient of the inter-division time with newborn size, not to be confused with the aforementioned correlation. Since the distribution is quite narrow, this is nearly the same as the correlation coefficient between logarithm of size and interdivision time, following the same logic which we used for the evaluation of the mother size:daughter size correlation coefficient.

Thus:

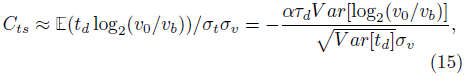

with *Var*[*t_d_*] the variance of the generation time which was calculated in the manuscript and found to be (Eq. (7) of main text):

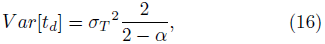

where *σ_T_* is the noise magnitude.

Similarly, the variance of the logarithm of newborn size,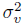, was found to be (Eq. (6) of main text):

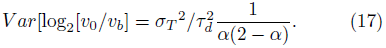

Combining these we find:

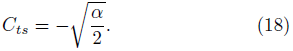

In particular, it is -1/2 for the incremental model.

### Reduction of biophysical model to the incremental model

We shall now show that the biophysical model of Ref. [2], where a constant volume is added between two DNA replication initiation events, perfectly realizes the incremental model (corresponding to Eq. (1) of the main), yet with Δ which depends on size in a particular way: Let us assume that that there are 2*^n^* replication forks at work (and hence 2*^n^***^+^**^1^ replication origins), and that initiation of DNA replication occurred at a volume *v*_i_ for one of them. Protein B will be accumulated at each origin until a critical amount is reached. This implies that the next initiation will occur (on average) at a volume 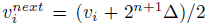, where according to the above biophysical mechanism Δ is *independent* of the growth rate. According to our assumption, there will be *n* division events from initiation to the end of replication. Since growth is exponential and we are assuming perfectly symmetric divisions, if the cell volume at initiation is **v*_i_* its volume at the end of DNA replication is *v^d^* = **v*_i_*2*^T_r_/τ_d_-n^*, regardless of when the intermediate division events happened. Its daughter cell will be born with a volume **v*_nb_* = *v^d^*/2, and its size at division will equal, by the same reasoning:

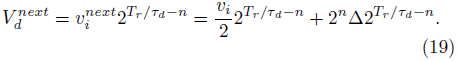

Thus we have (up to the noise):

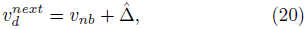

with:

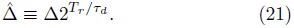

### Comparison of theory and numerics for time-additive noise

In this section we elaborate on the generality of the results derived in the main text by showing the excellent agreement between the analytical form of Eqs. (6) and (7) of the main text and the numerics. The distributions were evaluated numerically by following the lineage of 10^7^ divisions for each case.

Fig. 2 compares the analytical results for *α* = 1/2 with numerical results on the incremental model, for varying noise. We tested noise magnitudes ranging from the biologically relevant value of *σ*_*T*_/τ = 0.2 to relatively large noise with *σ*_*T*_/τ = 0.6. We find excellent agreement even for large noise - despite the fact that the incremental model is equivalent to the *α* = 1/2 analytically tractable model only to lowest order in (**v*_b_* – *v*_0_).

**FIG. 2.**
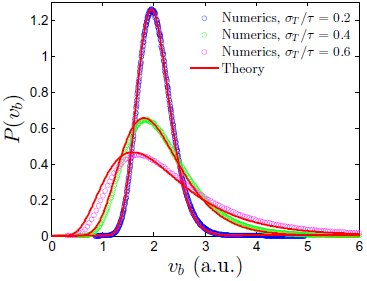
Numerical simulation compared with Eq. (6) of the main text, for the incremental model (corresponding to *α* = 1/2), for varying noise. The noise is added to the inter-division time and has a standard deviation *σ_T_*. For each case, the numerical distribution is extracted from a sequence of 10^7^ divisions.

### Solution of model for size-additive noise

Using the tools presented in the main text, it is straightforward to modify the approach for size-additive noise. In this case, it is convenient to use:

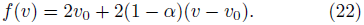

This agrees to lowest order in (*v* – *v*_0_) with the definition of *α* of the main text, and hence for the biologically relevant narrow distributions will be essentially equivalent. *α* = 0 has no size control, *α* = 1 realizes division at a constant size, and most importantly, *α* = 1/2 is an *exact* realization of the incremental model.

Now we assume that the noise is added to size rather than time, hence:

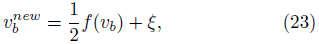

with ξ a Gaussian variable with standard deviation *σ_S_*. Using Eq. (27) we have:

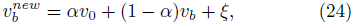

which can be rewritten as:

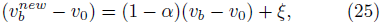

Following the same reasoning used in the main text, **v*_b_* – *v*_0_ will be a Gaussian variable with vanishing mean, and:

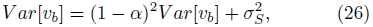

Hence:

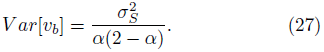

Fig. 3 verifies this result, comparing the analytic results for various values of *α* with numerics (including *α* = 1/2 which corresponds to the incremental model), all with size-additive noise with *σ*_*S*_/*v*_0_ = 0.27 (chosen to capture the width of the experimentally observed distribution).

**FIG. 3.**
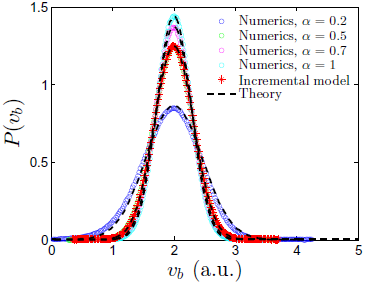
Comparison between the analytical results of a Gaussian distribution with variance given by Eq. (27) and numerics with *size-additive* noise, for varying values of *α*. Choosing *α* = 1/2 provides an exact realization of the incremental model.

### Distinguishing between size-additive and time-additive noise

For the realistic biological parameters, the difference between the time-additive and size-additive noise is not dramatic, as is shown in Fig. 4.

**FIG. 4.**
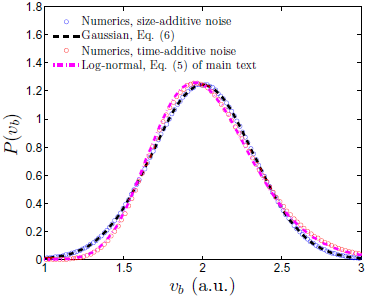
The distribution of newborn cell volume was found numerically, using the incremental model for both the cases of size-additive and time-additive noise. In the first case *σ*_*S*_/*v*_0_ = 0.2ln(2) ≈ 0.14, while in the latter *σ_T_*/*τ_d_* = 0.2. As is shown, the two cases are well-approximated by Gaussian and log-normal distributions, respectively, and by the theory corresponding to Eq. (27) of the SI and of Eq. (6) of the main text.

We also tested the size distributions resulting from the regulation strategy corresponding to a generic value of *α* (Eq. (3) in the main text), with a noise that has *both* a time-additive component (with standard deviation *σ_T_*) and a size-additive component (with standard deviation *σ_*S*_*). Based on the results described above that an approximately Gaussian distribution occurs for a size-additive noise, we expect that only the time-additive component of the noise will significantly contribute to the skewness of the distribution, since a Gaussian distribution has vanishing skewness. Therefore, we calculated numerically the skewness of the newborn size distribution: 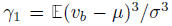, where *μ* ≈ *v*_0_ is the average newborn size and *σ* is the standard deviation of the size distribution.

As expected, we found that in order to have significant skewness, there has to be a time-additive noise component present, as is illustrated in Fig. 5.

**FIG. 5.**
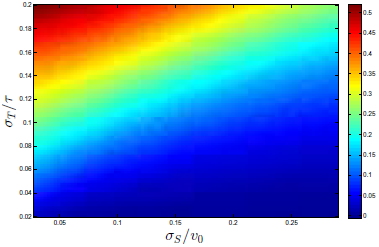
The skewness of the newborn size distribution is shown in the 2d map for the mixed case where both time-additive and size-additive noise are present, with magnitudes *σ_T_* and *σ_S_*. As is shown, distributions with non-negligible skewness only occur in the presence of time-additive noise.

Interestingly, in the analysis of financial markets a similar strategy is used to test the multiplicative nature of the stochastic processes underlying the market fluctuations [3, 4].

### Experimental support for time-additive noise

The purpose of this section is to show that existing experimental data for *E. coli* supports time-additive (i.e., multiplicative) noise rather than size-additive noise, using the tool developed in the previous section: namely, the skewness of the size distribution.

Ref. [5] measures the newborn size distribution of *E. coli* using the membrane elution technique, and shows that it has non-negligible positive skewness and it agrees very well with a log-normal distribution.

Recently, Ref. [1] analyzed cell size fluctuations and distributions. Their size distributions are strongly skewed to the right (γ_1_ > 0.5), suggesting that the noise is multiplicative rather than additive to the size. A similar analysis was performed in Ref. [6], which further corroborates our conclusion.

Although these various experiments suggest that noise in the size-control of bacteria is multiplicative rather than additive to size, a significant amount of additional experimental work is needed in order to elucidate the origin of the noise at the molecular level. Ref. [7] used the powerful tools of molecular biology to gain insights into precisely such a problem in budding yeast. In this pioneering work, they also model the noise as time-additive rather than size-additive, and are able to deduce the molecular mechanism for it and *modify it* in order to test their stochastic model. It would be highly rewarding to do the same for bacteria.

### DnaA as a volume integrator

Here, we review some recent progress in our understanding of the function of dnaA in initiating DNA replication in bacteria, and show that it shares many properties with the hypothetical model described in the main text. DnaA has two forms, an active, ATP-bound form, and an inactive ADP-bound form. It is believed that 20 dnaA proteins in their active form are needed in order to initiate DNA replication [8, 9]. This highly cooperative mechanism is parallel to the “critical number” of protein *B* in the main text (*P*_2_ in Ref. [2]). After initiation, the number of active copies of dnaA quickly drops [9], which is equivalent to the “degradation” of protein *B*. Importantly, dnaA is known to autoregulate [10], which in certain cases leads to a concentration approximately independent of the growth rate [11] - this was the requirement for protein *A* of the main text (*P*_1_ in Ref. [2]). As expected, inducing high levels of dnaA in *E. coli* leads to early initiation [12], while dnaA mutants initiate at a later time [13, 14]. Therefore, it is possible that the two forms of dnaA serve together to implement the volume integration which is hypothetically discussed in Ref. [2]. Certainly, many other proteins are involved in the process, such as SeqA [13], and further experiments are needed to better understand the biochemical and biophysical mechanisms. Nevertheless, the results presented in the main text lead to severe constraints on possible models.

## References

[1] S. Cooper, Bacterial growth and division: biochemistry and regulation of prokaryotic and eukaryotic division cycles (Elsevier, 1991).

[2] A. L. Koch, Bacterial growth and form (Springer, Berlin, 2001).

[3] P. Wang, L. Robert, J. Pelletier, W. L. Dang, F. Taddei, A. Wright, and S. Jun, Current Biology 20, 1099 (2010).

[4] J. Mnnik, F. Wu, F. J. H. Hol, P. Bisicchia, D. J. Sherratt, J. E. Keymer, and C. Dekker, PNAS (2012).

[5] Getting into shape: how do rod-like bacteria control their geometry?, A. Amir and S. van Teeffelen, invited review to appear in Systems and Synthetic Biology.

[6] A. L. Koch and M. Schaechter, Journal of General Microbiology 29, 435 (1962).

[7] M. Schaechter, J. P. Williamson, J. R. Hood, and A. L. Koch, Journal of General Microbiology 29, 421 (1962).

[8] P. Fantes, W. Grant, R. Pritchard, P. Sudbery, and A. Wheals, Journal of Theoretical Biology 50, 213 (1975).

[9] A. C. Chien, N. S. Hill, and P. A. Levin, Current biology 22, 340 (2012).

[10] A. C. Lloyd, Cell 154, 1194 (2013).

[11] P. Jorgensen and M. Tyers, Current Biology 14, R1014 (2004).

[12] D. O. Morgan, The Cell Cycle: Principles of Control (New Science Press, London, 2007).

[13] M. Osella, E. Nugent, and M. C. Lagomarsino, PNAS 111, 3431 (2014).

[14] L. Robert, M. Hoffmann, S. Krell, Nathalie Aymerich, J. Robert, and M. Doumic, BMC Biology 12, 17 (2014).

[15] W. Donachie, Nature 210, 1077 (1968).

[16] M. Schaechter, O. Maaløe, and N. Kjeldgaard, Journal of General Microbiology 19, 592 (1958).

[17] S. Cooper and C. E. Helmstetter, Journal of Molecular Biology 31, 519 (1968).

[18] L. J. Koppes, M. Meyer, H. B. Oonk, M. A. de Jong, and N. Nanninga, Journal of Bacteriology 143, 1241 (1980).

[19] L. Sompayrac and O. Maaløe, Nature 241, 133 (1973).

[20] W. Coffey, Y. P. Kalmykov, and J. T. Waldron, The Langevin Equation (World Scientific, Singapore, 1996).

[21] A. Amir, Y. Oreg, and Y. Imry, PNAS 109, 1850 (2012).

[22] R. N. Mantegna and H. E. Stanley, Introduction to econophysics: correlations and complexity in finance (Cambridge University Press, 2000).

[23] J.-P. Bouchaud and M. Potters, Theory of financial risks: from statistical physics to risk management, Vol. 12 (Cambridge University Press, Cambridge, 2000).

[24] S. Cooper, Theoretical Biology and Medical Modelling 3, 10 (2006).

[25] O. Sliusarenko, M. T. Cabeen, C. W. Wolgemuth, C. Jacobs-Wagner, and T. Emonet, PNAS 107, 10086 (2010).

[26] M. Godin, F. F. Delgado, S. Son, W. H. Grover, A. K. Bryan, A. Tzur, P. Jorgensen, K. Payer, A. D. Grossman, M. W. Kirschner, and S. R. Manalis, Nature methods 7, 387 (2010).

[27] M. Mir, Z. Wang, Z. Shen, M. Bednarz, R. Bashir, I. Golding, S. G. Prasanth, and G. Popescu, PNAS 108, 13124 (2011).

[28] M. Scott, C. W. Gunderson, E. M. Mateescu, Z. Zhang, and T. Hwa, Science 330, 1099 (2010).

[29] A. Amir and D. R. Nelson, PNAS 109, 9833 (2012).

[30] R. J. H. A. G. Marr and W. C. Trentini, Journal of Bacteriology 6, 91 (1966).

[31] See W. L. Voorn and A. J. H. Koppes, Arch Microbiol 169, 43 (1998) for a discussion on the history of the incremental model.

[32] S. D. Talia, J. M. Skotheim, J. M. Bean, E. D. Siggia, and F. R. Cross, Nature 448, 947 (2007).

[33] F. J. Trueba, O. M. Neijssel, and C. L. Woldringh, Journal of Bacteriology 150, 1048 (1982).

[34] G. Ullman, M. Wallden, E. G. Marklund, A. Mahmutovic, I. Razinkov, and J. Elf, Philosophical Transactions of the Royal Society B: Biological Sciences 368 (2013).

[35] F. Carrara, A. Giometto, A. Maritan, A. Rinaldo, F. Altermatt, et al., Proceedings of the National Academy of Sciences 110, 4646 (2013).

## References

[1] M. Osella, E. Nugent, and M. C. Lagomarsino, PNAS 111, 3431 (2014).

[2] L. Sompayrac and O. Maaløe, Nature 241, 133 (1973).

[3] R. N. Mantegna and H. E. Stanley, Introduction to econophysics: correlations and complexity in finance (Cambridge University Press, 2000).

[4] J.-P. Bouchaud and M. Potters, Theory of financial risks: from statistical physics to risk management, Vol. 12 (Cambridge University Press, Cambridge, 2000).

[5] L. J. Koppes, M. Meyer, H. B. Oonk, M. A. de Jong, and N. Nanninga, Journal of Bacteriology 143, 1241 (1980).

[6] L. Robert, M. Hoffmann, S. Krell, Nathalie Aymerich, J. Robert, and M. Doumic, BMC Biology 12, 17 (2014).

[7] S. D. Talia, J. M. Skotheim, J. M. Bean, E. D. Siggia, and F. R. Cross, Nature 448, 947 (2007).

[8] A. C. Chien, N. S. Hill, and P. A. Levin, Current biology 22, 340 (2012).

[9] W. D. Donachie and G. W. Blakely, Current Opinion in Microbiology 6, 146 (2003).

[10] R. E. Braun, K. O’Day, and A. Wright, Cell 40, 159 (1985).

[11] F. G. Hansen, T. Atlung, R. E. Braun, A. Wright, P. Hughes, and M. Kohiyama, Journal of Bacteriology 173, 5194 (1991).

[12] A. Löbner-Olesen, K. Skarstad, F. G. Hansen, K. von Meyenburg, and E. Boye, Cell 57, 881 (1989).

[13] E. Boye, T. Stokke, N. Kleckner, and K. Skarstad, Proceedings of the National Academy of Sciences 93, 12206 (1996).

[14] M. Lu, J. L. Campbell, E. Boye, and N. Kleckner, Cell 77, 413 (1994).

